# Myeloperoxidase activity predicts atherosclerotic plaque disruption and atherothrombosis

**DOI:** 10.1101/2023.10.08.561454

**Authors:** James Nadel, Xiaoying Wang, Prakash Saha, André Bongers, Sergey Tumanov, Nicola Giannotti, Weiyu Chen, Niv Vigder, Mohammed M. Chowdhury, Gastao Lima da Cruz, Carlos Velasco, Claudia Prieto, Andrew Jabbour, René M. Botnar, Roland Stocker, Alkystis Phinikaridou

## Abstract

**Background:** Unstable atherosclerotic plaque is characterized by increased myeloperoxidase (MPO) activity. As unstable plaque is vulnerable to disruption and ensuing thrombosis, we examined whether plaque MPO activity predicts atherothrombosis in a pre-clinical model and correlates with ruptured human atheroma.

**Methods:** To assess if plaque MPO activity predicts atherothrombosis, rabbits were subjected to aortic endothelial denudation, cholesterol feeding, *in vivo* magnetic resonance imaging (MRI) of MPO activity using MPO-Gd (gadolinium), followed by pharmacological triggering of atherothrombosis, histology, and MPO activity determined by liquid chromatography tandem mass spectrometry (LC-MSMS) by quantifying the MPO-specific product of hydroethidine, 2-chloroethidium. To correlate plaque MPO activity to ruptured human atheroma, *ex vivo* determination of MPO activity by MPO-Gd retention in carotid endarterectomy (CEA) specimens was correlated with *in vivo* MRI plaque phenotyping in patients, histology, and MPO activity determined by LC-MSMS.

**Results:** In rabbits, pre-trigger *in vivo* MPO activity, validated by LC-MSMS and histology, was higher in thrombosis-prone than thrombosis-resistant plaques and lesion-free segments (R1 relaxation rate = 2.2 ± 0.4 versus 1.6 ± 0.2 and 1.5 ± 0.2 s^-1^, respectively, p<0.0001), and it predicted atherothrombosis. In CEA specimens, MPO-Gd retention was greater in histologic and MRI-graded American Heart Association (AHA) type VI than types III, IV and V plaques (ΔR1 relaxation rate from baseline = 48 ± 6 versus 16 ± 7, 17 ± 8 and 23 ± 8%, respectively, p<0.0001). This association was confirmed by comparing AHA grade to MPO activity determined by LC-MSMS (277 ± 338 versus 7 ± 6, 11 ± 12 and 42 ± 39 pmol 2-chloroethidium/mg protein for type VI versus type III-V plaques, respectively, p=0.0008).

**Conclusions:** MPO activity is elevated in thrombosis-prone rabbit and ruptured human atheroma. Non-invasive molecular imaging of MPO activity predicts atherothrombosis, highlighting the potential of arterial MPO activity to detect vulnerable, destabilized atherosclerosis.

## Introduction

To reduce cardiovascular morbidity and mortality, clinical research interest has turned to investigating imaging techniques that reliably detect atherosclerotic plaques prone to thrombosis. Such advancements could improve patient care and reduce the healthcare burden by aiding decision-making for therapeutic intervention for those with ‘vulnerable’ plaques.^1^ Currently, intervention is based on the severity of luminal stenosis, even though it is well established that stenosis alone is an inconsistent predictor of cardiovascular events.^2–6^ One reason for the continued reliance on stenotic grade is the lack of available non-invasive strategies for the identification of vulnerable and destabilized plaques.

Current non-invasive imaging strategies infer plaque vulnerability from structural determinants of plaque composition or non-specific features of arterial inflammation.^7^ Magnetic resonance imaging (MRI) is a powerful tool for imaging carotid and large vessel atherosclerosis due to its high spatial resolution and soft-tissue contrast^8^ and it has excellent correlation with histology, permitting delineation and quantification of plaque components responsible for plaque vulnerability.^9–11^ Nonetheless, available MRI techniques fail to determine pathobiological activity of plaques, meaning only surrogate markers of plaque destabilization can be garnered.^12^ The development of molecular probes that provide insight into plaque pathological activity beyond plaque composition, burden and luminal stenosis may improve stratification and management of patients with atherosclerotic diseases.^13^

Myeloperoxidase (MPO) is a pro-inflammatory enzyme that performs a unique role within the innate immune response, converting hydrogen peroxide and chloride to microbicidal hypochlorous acid within the phagolysosome.^14^ Despite these protective properties, up to 30% of neutrophil MPO can be released into the extracellular space and produce hypochlorous acid causing tissue injury, including atherosclerotic plaque destabilization.^15–18^ A gadolinium-based MRI probe (gadolinium-*bis*-5-hydroxytryptamide diethylenetriaminepentaacetic acid, MPO-Gd) identifies extracellular MPO activity in atherosclerotic plaques in a rabbit model.^19^ MPO-mediated oxidation of the phenol moiety of 5-hydroxytryptamide forms MPO-Gd radicals that undergo oligomerization and protein-probe adduct formation.^20^ This results in enhanced retention of gadolinium in tissues with increased MPO activity and shortening of T1 relaxation time, giving rise to enhanced signal in T1-weighted MRI.^19, 21, 22^ Recently MPO-Gd MRI was shown to accurately detect histologically unstable plaque in mouse models of plaque instability.^23^ Moreover, in the *Apoe*^-/-^ mouse model of tandem stenosis, deletion of the *Mpo* gene decreased formation of vulnerable plaque,^23^ and pharmacological inhibition of MPO activity stabilized pre-existing unstable plaque.^24^ These findings highlight the causal role of extracellular MPO activity in plaque destabilization and confirm MPO-Gd’s utility to image MPO activity in plaque.

However, the capacity of plaque MPO activity as a non-invasive risk predictor of plaque disruption and thrombosis has not been examined. We hypothesized that increased plaque MPO activity is predictive of plaque destabilization and ensuing thrombosis, and that MPO-Gd enables non-invasive detection of vulnerable/culprit atherosclerotic lesions. We tested this hypothesis by assessing MPO activity using MPO-Gd and a previously validated method based on liquid chromatography tandem mass spectrometry (LC-MSMS)^25, 26^ in an animal model of atherothrombosis and in ruptured and destabilized human plaques.

## Methods

### Data Availability

Detailed Materials and Methods, Major Resources Table, and additional data supporting the findings of this study are available in the **Supplemental Material**. Raw data that support the findings of this study are available from the corresponding authors upon reasonable request.

## Results

To assess whether plaque MPO activity predicts atherosclerotic plaque disruption, we used a robust model of trigger-induced atherothrombosis in rabbits^27^ (n = 12) and employed non-invasive *in vivo* molecular imaging of MPO activity using MPO-Gd in combination with *ex vivo* determination of MPO activity by LC-MSMS by quantifying the MPO-specific product of hydroethidine, 2-chloroethidium.

### Molecular Imaging of Arterial MPO Activity Predicts Trigger-Induced Atherothrombosis

All twelve rabbits developed atherosclerosis, and five advanced to trigger-induced atherothrombosis. A total of 236 aortic segments were analysed pre-trigger using 2D T1-weighted black blood (T1BB), 3D inversion recovery T1w (T1w-IR) and 3D T1 mapping images. These segments represented lesion-free artery (n = 87) and plaques (n = 149) of various disease stages and trigger-induced outcomes (**Figures S1B-C & S2**). Of the 149 plaque-containing segments, 24 (16%) developed trigger-induced thrombosis with the remaining 125 (84%) categorized as thrombosis-resistant, with stable plaques.

Representative *in vivo* rabbit MRI images from the three groups of aortic segments acquired at week 12 are shown in **Figure 1**, with the corresponding images at week 8 shown in **Figure S3**. Prior to MPO-Gd administration, T1w-IR images showed very low aortic signal intensity (**Figure 1A**), with comparable R1-values for the aortic wall and paraspinal muscle (**Figure 1B, blue color**). Late MPO-Gd enhancement of patchy appearance was observed in all three groups, although plaque-containing segments had higher signal intensity and R1-values compared with lesion-free segments.

**Figure 1.**
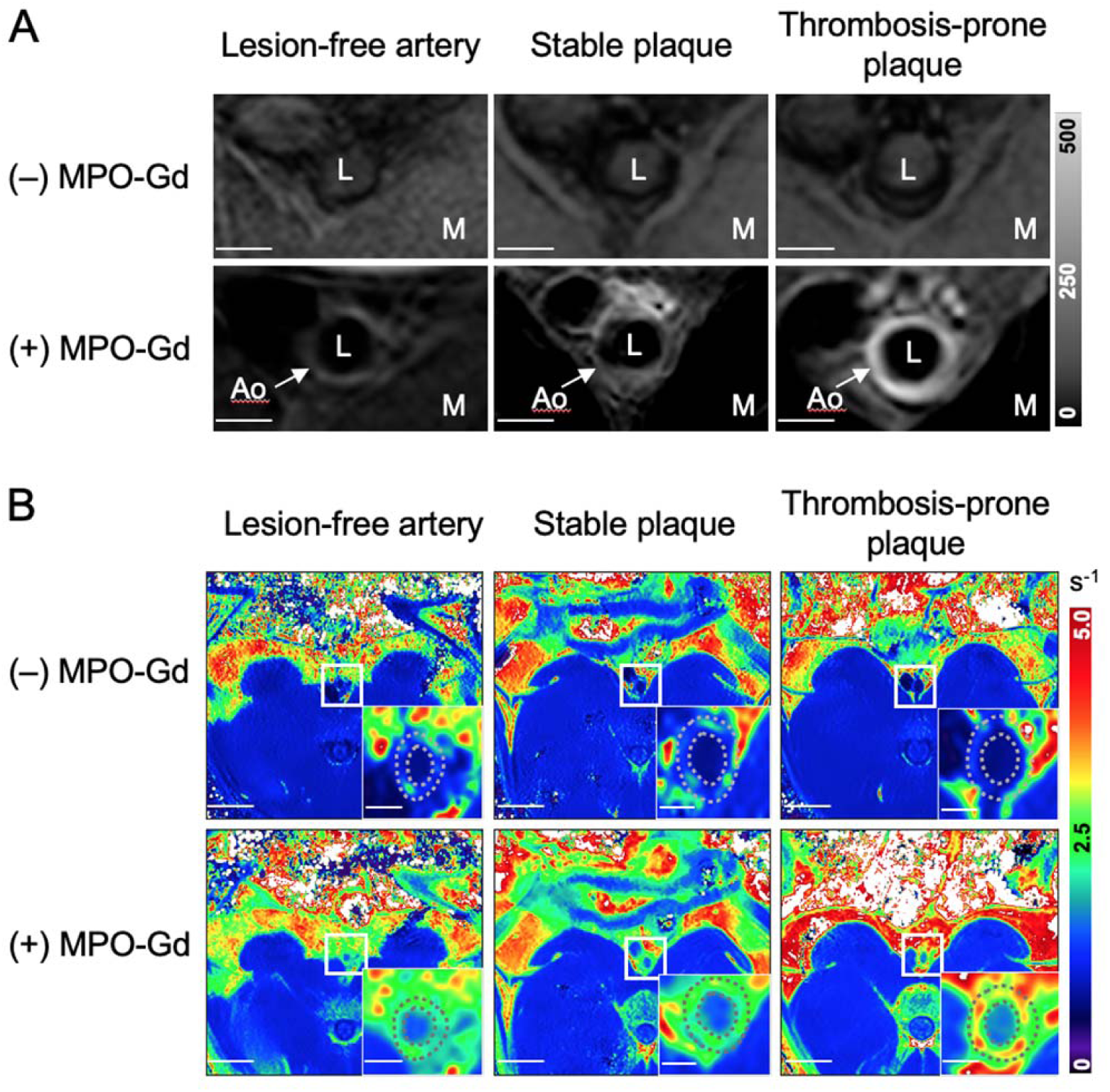
*In vivo* pre-trigger MR images of lesion-free abdominal aortic segments and segments containing stable or thrombosis-prone plaque in rabbits 12 weeks after commencement of cholesterol-fortified diet. **A,** 2D fat suppressed quadruple inversion recovery black blood T1 weighted turbo spin echo (T1w-IR) images before (–) and 1 h after (+) injection of MPO-Gd. Scale bar 2 mm. **B,** R1 relaxation maps before (–) and 1.5 h after (+) injection of MPO-Gd. Scale bars are 20 and 2 mm for main panels and insets, respectively. Ao, aortic wall; L, aortic lumen; M, paraspinal muscle.

Quantitative MRI analysis of the week 12 pre-trigger images showed similar aortic wall areas (p > 0.9) measured on T1BB images and late MPO-Gd-enhanced area (p = 0.2) measured on T1w-IR images in stable and thrombosis-prone plaques (**Figure 2A-B**). However, thrombosis-prone plaques had significantly higher aortic tissue-to-muscle contrast ratio (9.4 ± 6.6 versus 3.2 ± 2.0) and R1-values (2.2 ± 0.2 versus 1.6 ± 0.2 s^-1^) at week 12 compared with stable plaques (p < 0.0001 for both comparisons) **(Figure 2C-D**). Comparison of lesion-free aortic segments and segments with stable plaque at 12 weeks showed no difference in the aortic tissue-to-muscle contrast ratios (p = 0.4) and R1-values (p = 0.5). Comparison with the corresponding quantitative MRI analyses for week 8 MRI data showed overall similar trends (**Table S1**).

**Figure 2.**
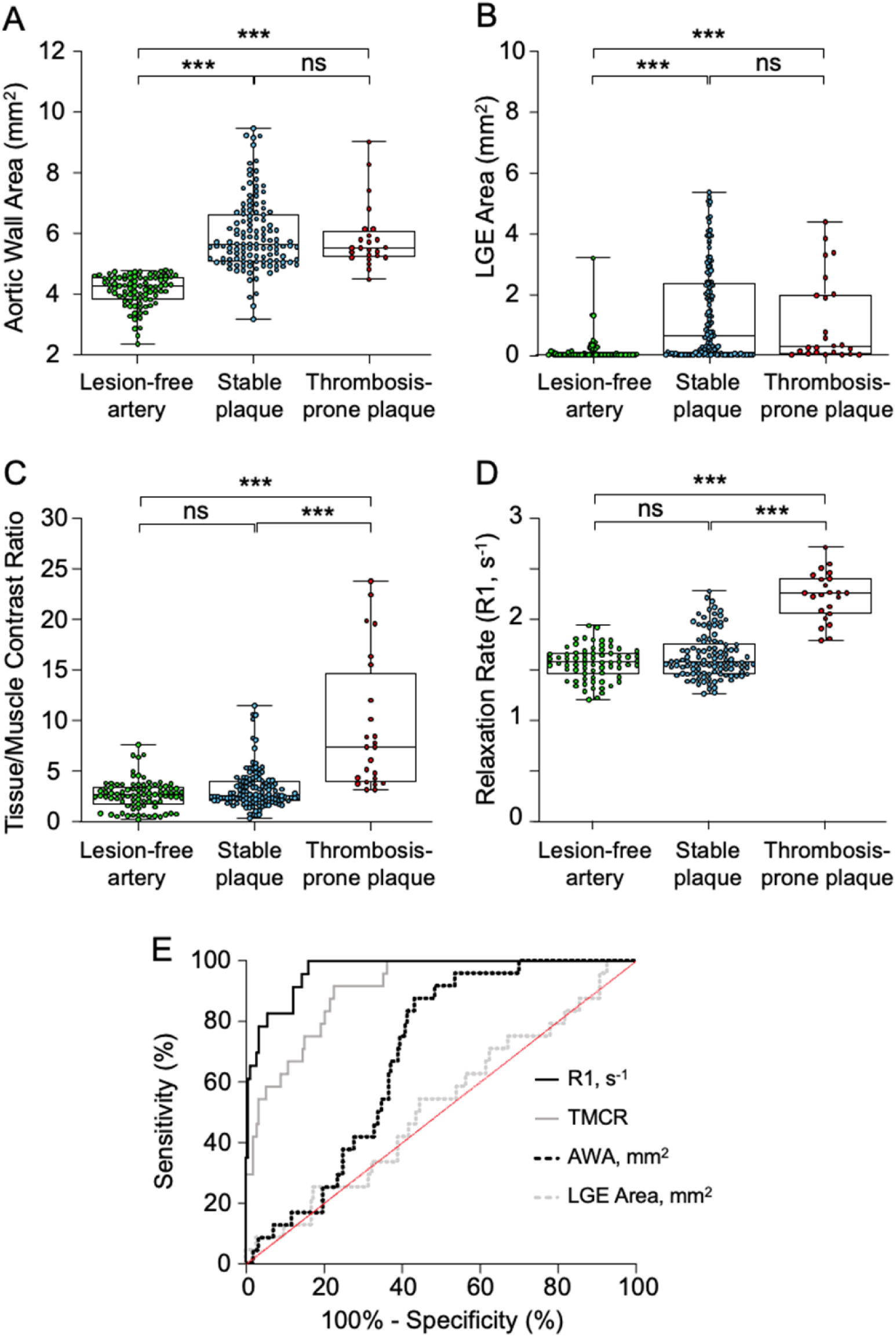
Quantitative analysis of MPO-Gd enhanced *in vivo* pre-trigger MR images of lesion-free segments and segments containing stable or thrombosis-prone plaque in rabbits 12 weeks after commencement of cholesterol-fortified diet. **A,** Aortic wall area (AWA) segmented from T1BB images. **B,** Late MPO-Gd-enhanced (LGE) area on masked 2D fat suppressed quadruple inversion recovery black blood T1 weighted turbo spin echo (T1w-IR) images 1 h after MPO-Gd administration. **C,** LGE signal intensity presented as tissue-to-muscle contrast ratio (TMCR) on masked T1w-IR images 1 h after MPO-Gd administration. **D,** R1 relaxation rate on masked T1 maps 1.5 h after MPO-Gd administration. **E,** Receiver operating characteristic (ROC) curves of AWA, LGE area, TMCR, and R1 in predicting trigger-induced thrombosis. ***p <0.001.

The ROC curves showed that the pre-trigger R1-value was the best and aortic tissue-to-muscle contrast ratio a good predictor for trigger-induced thrombosis with high sensitivity (both 100%) and specificity at both time points (86% and 84%, respectively) after administration of MPO-Gd (**Figure 2E**). By comparison, aortic wall area and late MPO-Gd-enhanced area were poor indicators for trigger-induced thrombosis at both time points (**Figure S4**). **Table S1** provides a more detailed analysis, including confidence intervals and cut-offs with corresponding sensitivities and specificities.

### Increased MPO Activity in Thrombosis-Prone Rabbit Plaques in the Absence of Differences in Plasma MPO Protein Concentrations

Next the results obtained with *in vivo* molecular MRI using MPO-Gd were validated by *ex vivo* quantifying the MPO-specific product 2-chloroethidium (2-Cl-E^+^) from added hydroethidine by LC-MSMS. MPO activity was significantly higher in thrombosis-prone plaques compared with stable plaques (80.4 ± 30.0 versus 20.9 ± 8.4 pmol/mgp, p = 0.004) and lesion-free aortic segments (20.7 ± 6.9 pmol/mgp, p=0.02) (**Figure 3A**). Moreover, MPO activity determined by LC-MSMS showed a positive correlation with R1-values determined from the *in vivo* T1 maps (r = 0.62) (**Figure 3B**). Finally, IHC analysis (**Figure 3C**) showed that MPO protein was more abundant in thrombosis-prone than stable plaques, and rarely detected in the lesion-free aortic sections. Areas of high MPO expression correlated with the spatial distribution of MPO-Gd enhancement seen on pre-trigger T1w-IR images. Venous blood samples collected from rabbits at 8 and 12 weeks before triggering of atherothrombosis showed no difference in the plasma MPO protein concentration between animals that developed trigger-induced atherothrombosis and those with stable plaque (8 weeks: 3.1 ± 0.4 versus 3.5 ± 0.4 ng/mL, p = 0.10, and 12 weeks: 2.8 ± 1.0 versus 3.3 ± 0.8, p = 0.47, respectively) (**Figure S8A).**

**Figure 3.**
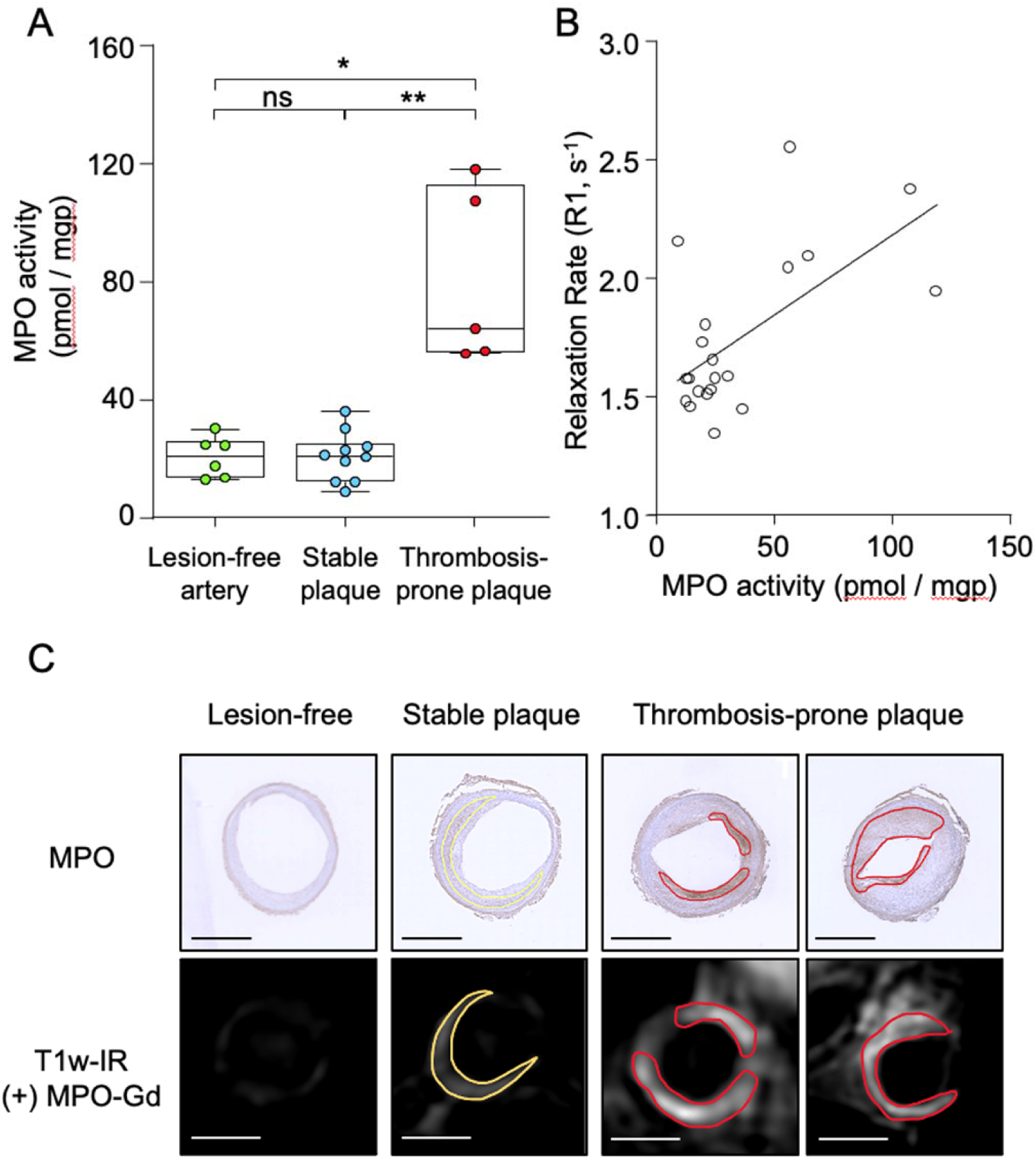
Correlation of *in vivo* MRI data with MPO activity determined by liquid chromatography tandem mass spectrometry in lesion-free rabbit aortic segments and segments containing stable or thrombosis-prone plaque. **A,** Aortic segments containing thrombosis-prone plaques have higher protein-standardized MPO activity expressed as pmol 2-chloroethidium per mg of protein (2-Cl-E^+^/mgp) than the segments containing stable plaques and lesion-free segments. **B,** Correlation between arterial MPO activity and R1-values. **C,** Co-localization of MPO protein assessed by immunohistochemistry and late MPO-Gd enhancement seen on MRI images. Scale bars 2 mm. *p <0.05, **p <0.01.

### MPO-Gd Activity as a Marker of Ruptured Plaques in Humans

Demographics and characteristics of the clinical study population are presented in **Table 2**. Patients were predominantly older Caucasian males on optimal medical therapy, with lipid profiles within guideline-directed ranges. Forty percent of the cohort had symptomatic carotid disease with 30% having a confirmed ipsilateral stroke on pre-surgical neuroimaging. *In vivo* carotid MRI data is presented in **Table 3**. 12 out of 30 patients underwent dedicated carotid MRI for plaque characterization prior to CEA (**Figure S5**), with most having severe carotid stenoses by NASCET criteria. Most slices analysed by MRI were mature AHA type V lesions, and 17% had features of AHA type VI ruptured and destabilized plaques (see **Table 3** for details). Interobserver discordance occurred with 24 out of the 101 slices assigned an MRI-based AHA grade.

**Table 1.** ROC Analysis for Imaging Indexes Predicting Trigger-Induced Atherothrombosis in Rabbits.

|  | **Aortic wall area**<br>**(mm<sup>2</sup>)** | **Late MPO-Gd**<br>**enhanced area**<br>**(mm<sup>2</sup>)** | **Tissue-to-Muscle**<br>**Contrast Ratio** | **R1 (s<sup>-1</sup>)** |
| --- | --- | --- | --- | --- |
| **Week 8** |  |  |  |  |
| Area Under Curve | 0.66 | 0.59 | 0.88 | 0.96 |
| 95% CI | 0.57-0.76 | 0.47-0.72 | 0.82-0.94 | 0.93-0.98 |
| p-value | 0.009 | 0.142 | <0.0001 | <0.0001 |
| Cut-off | > 5.69 mm<sup>2</sup> | > 0.04 mm<sup>2</sup> | > 2.83 | > 1.77 |
| Sensitivity, % | 54.2 | 66.7 | 87.5 | 100.0 |
| Specificity, % | 71.7 | 40.1 | 75.9 | 86.1 |
| **Week 12** |  |  |  |  |
| Area Under Curve | 0.69 | 0.52 | 0.91 | 0.97 |
| 95% CI | 0.61-0.77 | 0.40-0.64 | 0.86-0.96 | 0.94-0.99 |
| p-value | 0.003 | 0.796 | <0.0001 | <0.0001 |
| Cut-off | > 5.40 mm<sup>2</sup> | > 0.04 mm<sup>2</sup> | > 3.69 | > 1.82 |
| Sensitivity, % | 62.5 | 58.3 | 91.7 | 100.0 |
| Specificity, % | 63.2 | 43.4 | 77.4 | 83.9 |

**Table 2.**
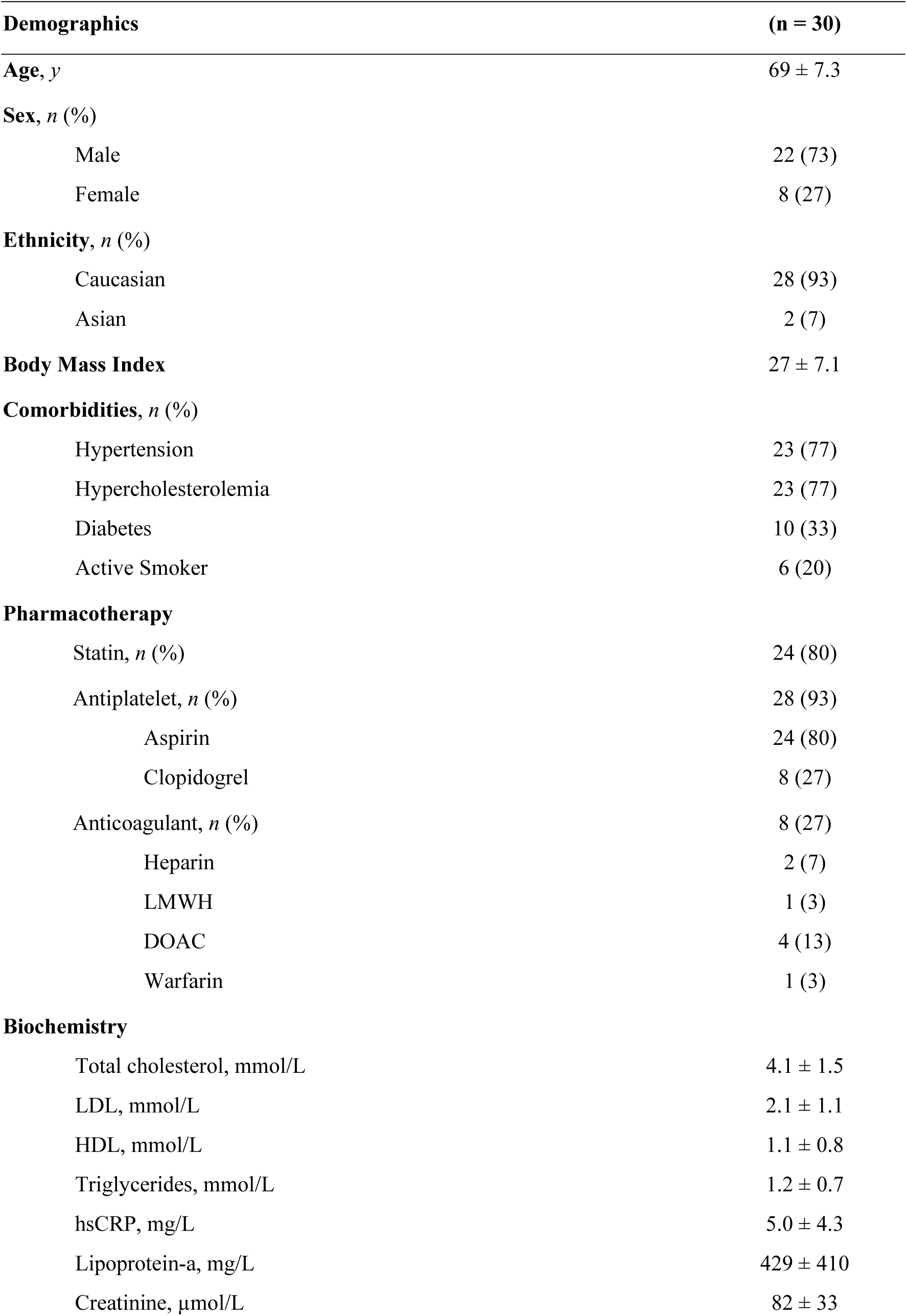
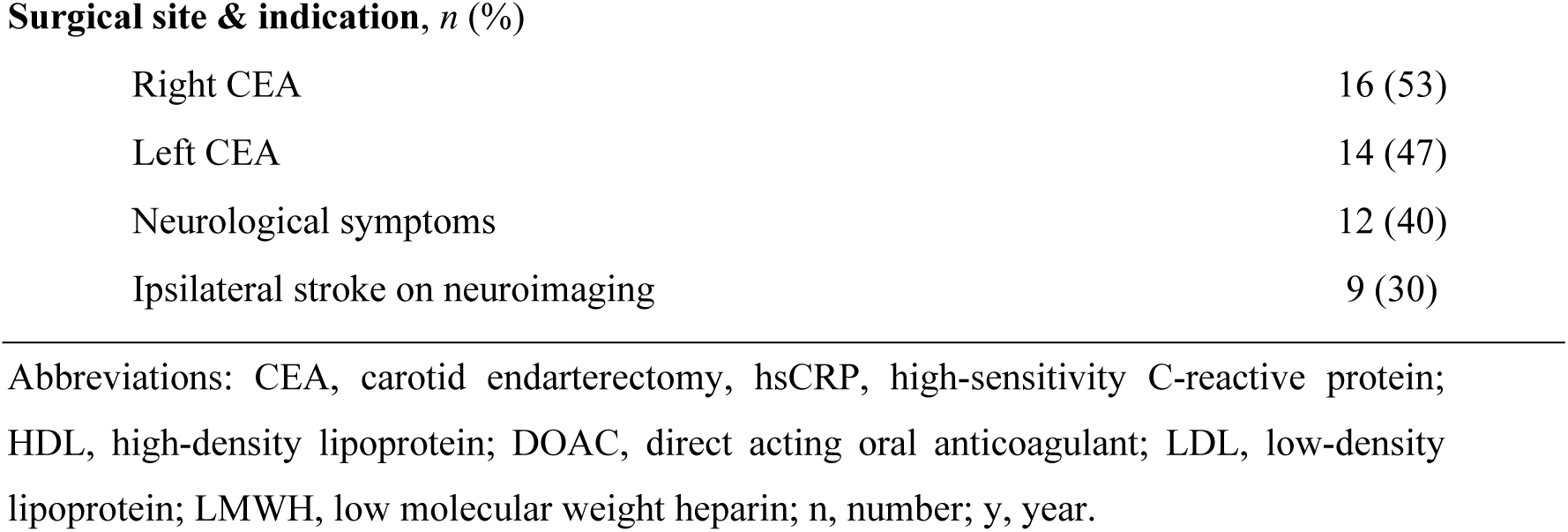
Cohort characteristics.

| **Demographics** | **(n = 30)** |
| --- | --- |
| **Age, y** | 69 ± 7.3 |
| **Sex, n (%)** |  |
| Male | 22 (73) |
| Female | 8 (27) |
| **Ethnicity, n (%)** |  |
| Caucasian | 28 (93) |
| Asian | 2 (7) |
| **Body Mass Index** | 27 ± 7.1 |
| **Comorbidities, n (%)** |  |
| Hypertension | 23 (77) |
| Hypercholesterolemia | 23 (77) |
| Diabetes | 10 (33) |
| Active Smoker | 6 (20) |
| **Pharmacotherapy** |  |
| Statin, n (%) | 24 (80) |
| Antiplatelet, n (%) | 28 (93) |
| Aspirin | 24 (80) |
| Clopidogrel | 8 (27) |
| Anticoagulant, n (%) | 8 (27) |
| Heparin | 2 (7) |
| LMWH | 1 (3) |
| DOAC | 4 (13) |
| Warfarin | 1 (3) |
| **Biochemistry** |  |
| Total cholesterol, mmol/L | 4.1 ± 1.5 |
| LDL, mmol/L | 2.1 ± 1.1 |
| HDL, mmol/L | 1.1 ± 0.8 |
| Triglycerides, mmol/L | 1.2 ± 0.7 |
| hsCRP, mg/L | 5.0 ± 4.3 |
| Lipoprotein-a, mg/L | 429 ± 410 |
| Creatinine, μmol/L | 82 ± 33 |
| **Surgical site & indication, *n* (%)** |  |
| Right CEA | 16 (53) |
| Left CEA | 14 (47) |
| Neurological symptoms | 12 (40) |
| Ipsilateral stroke on neuroimaging | 9 (30) |

Abbreviations: CEA, carotid endarterectomy; hsCRP, high-sensitivity C-reactive protein; HDL, high-density lipoprotein; DOAC, direct acting oral anticoagulant; LDL, low-density lipoprotein; LMWH, low molecular weight heparin; n, number; y, year.

**Table 3.** *In vivo* MRI for Patient.

| ***In vivo* carotid MRI** |  |  |
| --- | --- | --- |
| **Analysis, *n* (%)** |  |  |
| Scans performed |  | 12/30 (40%) |
| Total slices analysed |  | 101 |
| Average slices analysed per patient |  | 8.4 |
| **MRI-based AHA grade, *n* (%)** |  |  |
| III |  | 24 (24) |
| IV |  | 17 (17) |
| V |  | 43 (43) |
| VI |  | 17 (17) |
| **Plaque features** |  |  |
| Luminal stenosis, % |  |  |
| NASCET criteria |  | 77 ± 4.0 |
| ESCT criteria |  | 82 ± 5.4 |
| Total plaque volume, cm<sup>3</sup> |  | 0.6 ± 0.8 |
| Total lipid volume, cm<sup>3</sup> |  | 0.08 ± 0.1 |
| Total lipid : total plaque volume |  | 0.13 ± 0.1 |
| Fibrous cap thinning/loss, *n* (%) |  | 4 (33) |
| Intraplaque hemorrhage, *n* (%) |  | 4 (33) |
| Thrombus, *n* (%) |  | 2 (17) |
| **Clinical disposition, *n* (%)** |  |  |
| Neurological symptoms |  | 5 (42) |
| Ipsilateral stroke on neuroimaging |  | 6 (50) |

Abbreviations: ESCT, European Carotid Surgery Trial; NASCET, North American
Symptomatic Carotid Endarterectomy Trial; n; number.

Following optimization of the *ex vivo* molecular MRI method using MPO-Gd (**Supplemental Material**), R1 relaxation rates relative to pre-contrast baseline values were deemed to be a reliable surrogate of specific MPO-Gd retention in CEA sections 4 h after probe activation (**Figure S7**). First T1 mapping sequences reflecting MPO-Gd retention/MPO activity were compared with histological features (**Figure 4A-B**). Areas of MPO-Gd retention co-localized to immunohistochemical MPO staining, particularly in the shoulder regions of plaques. MPO activity was higher in CEA specimens that contained ruptured compared with stable plaques, as determined by the R1 relaxation rate (1.3 ± 0.1 versus 0.8 ± 0.2 s^-1^, p <0.0001) and ΔR1 values from baseline (49 ± 4 versus 17 ± 9 %, p <0.0001) (**Figure 4C-D**). Similarly, when comparing *ex vivo* MPO-Gd retention in whole CEA samples to *in vivo* MRI-determined AHA grade, probe retention corresponded to regions of MPRAGE hyperintensity and cap disruption in slices containing destabilized plaques (**Figure 5A-B**). Following MPO-Gd activation, ruptured/destabilized type VI plaques had higher mean R1-values (1.2 ± 0.1 s^-1^) than types III, IV and V plaques (0.8 ± 0.1, 0.8 ± 0.1 and 0.8 ± 0.2 s^-1^, respectively) (p <0.0001) (**Figure 5C**). Similarly, type VI plaques had higher ΔR1 (48 ± 6 %) than types III - V plaques (16 ± 7, 17 ± 8 and 23 ± 8 %, respectively) (p <0.0001) (**Figure 5D**), indicative of increased MPO activity in ruptured/destabilized plaques.

**Figure 4.**
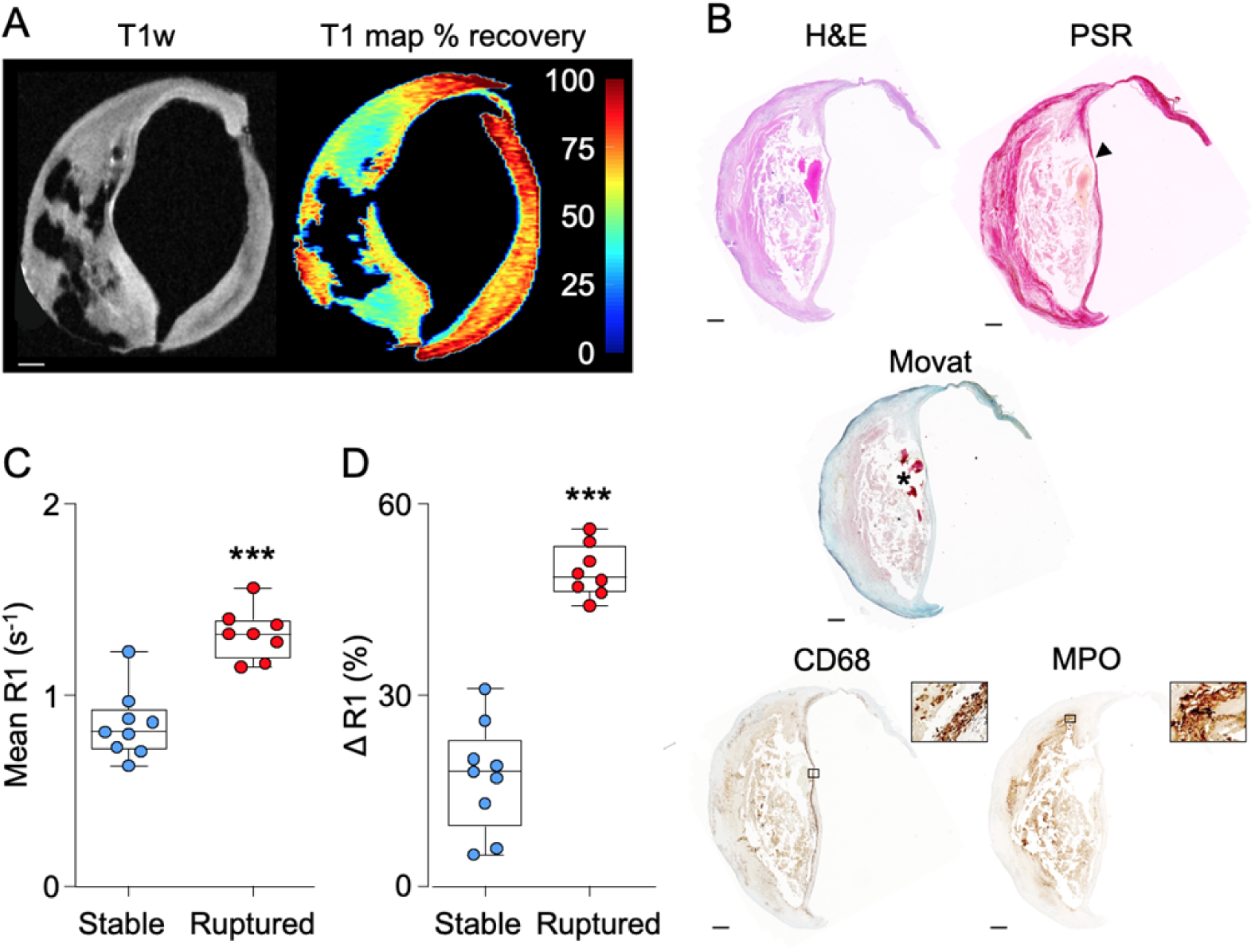
Comparison between MPO activity assessed by *ex vivo* MPO-Gd retention and AHA grading determined by histology. **A,** Baseline T1w *ex vivo* MR image with corresponding T1 colored map of percentage recovery from baseline following MPO-Gd probe activation. Scale bar 1 mm. **B,** Corresponding histological section showing a thin-capped (arrowhead) fibroatheroma with underlying intraplaque hemorrhage (asterisk) as well as MPO and CD68 positive staining (magnified boxes x 100). Scale bar 500 μm. Areas of MPO-Gd probe retention correspond to MPO IHC particularly in the shoulder regions of the plaque. **C-D,** Box and whisker plots showing relaxation rate (R1, s^-1^) and ΔR1 (%) following MPO-Gd activation in stable and ruptured plaques. Ruptured plaques have higher relaxation rate R1 (1.3 ± 0.1 versus 0.8 ± 0.2 s^-1^, p <0.0001) and ΔR1 (49 ± 4 versus 17 ± 9 %, p <0.0001) than stable plaques. ***p <0.001.

**Figure 5.**
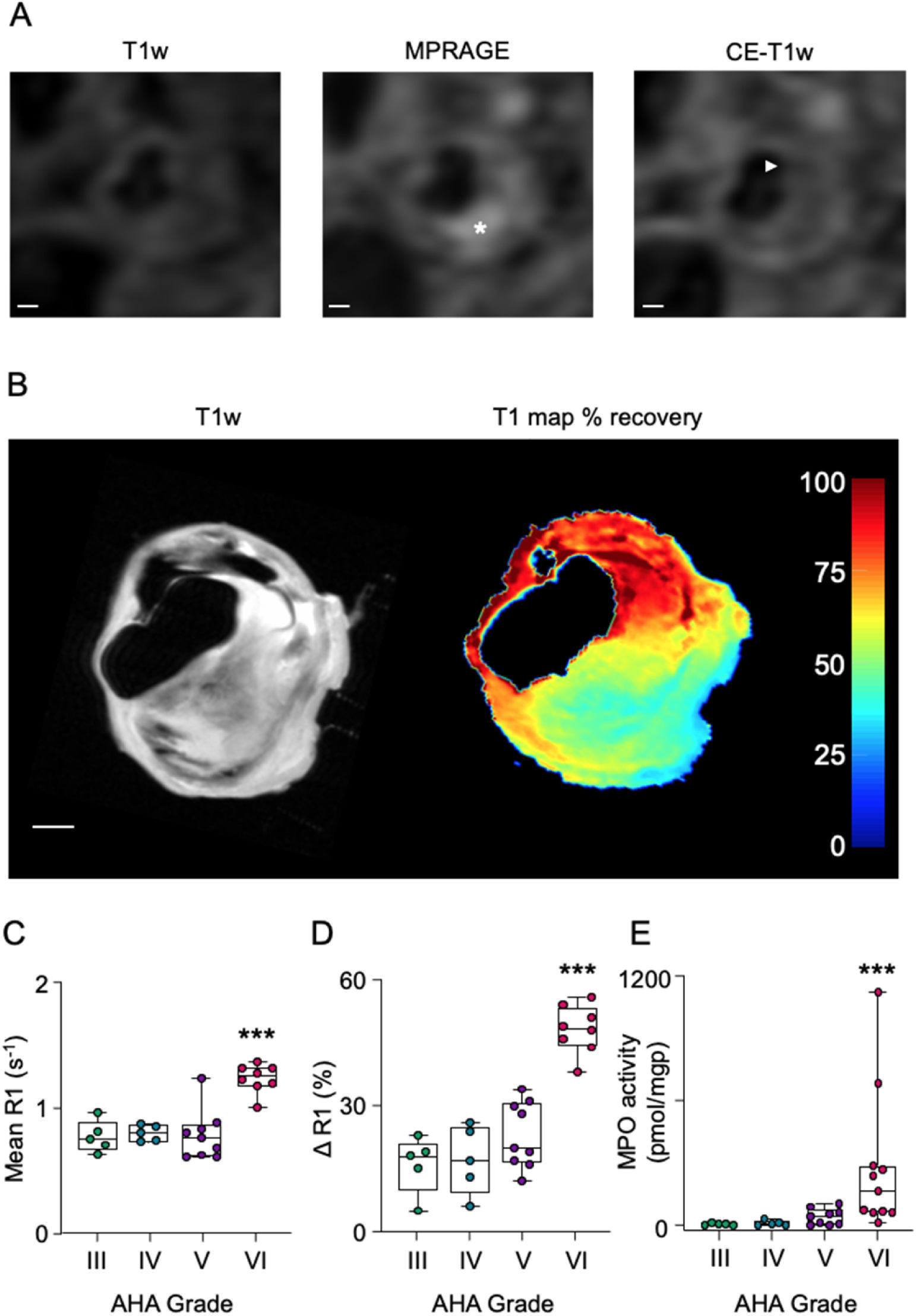
Comparison between AHA grading determined by *in vivo* MRI and MPO activity assessed by *ex vivo* MPO-Gd retention and determination of the conversion of hydroethidine to 2-chloroethidium (2-Cl-E^+^) by liquid chromatography tandem mass spectrometry. **A,** *In vivo* carotid MRI showing a lipid-rich plaque with intraplaque hemorrhage (high intensity, asterisks) on the MPRAGE sequence and fibrous cap thinning on CE T1w (arrowhead) images. **B,** T1 *ex vivo* MRI with T1 colored map of percentage recovery from baseline following MPO-Gd probe activation. Probe retention corresponds with the region of MPRAGE hyperintensity on *in vivo* imaging. Scale bars 1 mm. **C-D,** Box and whisker plots showing mean relaxation rate R1 and ΔR1 following MPO-Gd activation in plaques graded by MRI-based AHA classification. Following MPO-Gd activation, ruptured/destabilized type VI plaques have higher mean R1 (1.2 ± 0.1 s^-1^) than types III - V plaques (0.8 ± 0.1, 0.8 ± 0.1 and 0.8 ± 0.2 s^-1^, respectively) (p <0.0001). Similarly, type VI plaques have higher ΔR1 (48 ± 6 %) than types III - V plaques (16 ± 7, 17 ± 8 and 23 ± 8 %, respectively) (p <0.0001). **E,** Type VI plaques determined histologically have higher MPO activity, expressed as pmol 2-Cl-E^+^ per mg of protein than type III-V plaques (277 ± 338 pmol/mgp versus 7 ± 6, 11 ± 12 and 42 ± 39 pmol/mgp, respectively; p = 0.0008). ***p <0.001.

We next compared *ex vivo* MRI data with histologic and MRI-based plaque assessment to internally validate MRI-based analyses. MPO-Gd retention was similar for histologically stable (ΔR1 = 17 ± 9 %, n = 9) and MRI-graded types III-V (ΔR1 = 20 ± 8 %, n = 19) plaques (p = 0.8), and for histologically ruptured (49 ± 4 %, n = 8) and MRI-graded type VI (48 ± 6 %, n = 8) plaques (p = 0.6).

The validity of MPO-Gd to detect MPO activity in CEA sections was confirmed through the quantification of the MPO specific adduct 2-Cl-E^+^ in an *ex vivo* reaction using LC-MSMS. MPO activity was significantly higher in type VI plaques compared with histologic defined type III-V plaques (277 ± 338 pmol/mgp versus 7 ± 6, 11 ± 12 and 42 ± 39 pmol/mgp, respectively; p = 0.0008) (**Figure 5E**). Lastly, blood taken prior to CEA surgery showed an increase in the mean plasma MPO protein concentration in patients with ruptured plaques compared to those with stable plaques (268 ± 136 versus 179 ± 62 ng/mL; p = 0.02) (**Figure S8B**).

## Discussion

The present study sought to determine the relationship between plaque MPO activity and rupture and the utility of measuring plaque MPO activity for predicting atherothrombosis. The results presented establish for the first time that elevated arterial MPO activity detected *in vivo* by molecular MRI employing MPO-Gd discriminates between thrombosis-prone and stable plaques and predicts future plaque disruption in a rabbit model of atherothrombosis. The data also demonstrate that MPO activity, as measured by quantitative R1 relaxation rates and confirmed by LC-MSMS analysis, is increased in ruptured human atherosclerotic plaques compared with stable plaques. These results highlight that arterial MPO activity detects culprit lesions and can predict future susceptibility to atherothrombosis. They identify molecular imaging of MPO activity as a promising non-invasive strategy for the detection of vulnerable plaque.

Our results suggest that non-invasive imaging of plaque MPO activity is a promising candidate for clinical translation to predict adverse prognosis and guide treatment. Indeed, in the rabbit model of atherothrombosis used, arterial MPO activity represented by R1 relaxation rates positively correlated with MPO activity measured by LC-MSMS, and R1 relaxation values reliably predicted trigger-induced plaque disruption with high sensitivity. IHC results showed that MPO protein co-localized with sites of high MPO activity (detected by *in vivo* MPO-Gd) and thrombosis-prone plaque. Strikingly, MPO activity was significantly elevated in thrombosis-prone plaques as early as 8 weeks and remained persistently high at 12 weeks, while the plasma concentrations of MPO remained comparable at both time points. This suggests that MPO activity may be an early pre-thrombosis diagnostic marker that precedes changes in circulating MPO protein and the advanced plaque features currently relied upon by clinical MRI sequences including intraplaque hemorrhage (IPH) and cap thinning/disruption.

While in the rabbit model plaque disruption was ‘triggered’ using Russell’s viper venom as a procoagulant factor and histamine as a vasopressor and endothelial toxin, plaque disruption depends on an unstable plaque phenotype, thereby resembling atherothrombosis in humans.^27, 28^ The model also offers the possibility of imaging plaque activity at precise time points prior to ‘triggering’ atherothrombosis, making it a unique tool to develop and test imaging strategies for the detection of unstable plaque prior to cardiovascular events. As plaque disruption is caused by both plaque rupture and erosion,^27^ the rabbit study does not allow differentiation between plaque rupture and erosion as the cause of trigger-induced thrombosis. The role of plaque MPO activity in plaque disruption and its potential to predict susceptibility to atherothrombosis is supported by a recent study reporting elevated MPO activity in carotid plaque to correlate with symptomatic carotid disease and stroke.^18^

Three separate lines of evidence support the notion of increased MPO activity in human ruptured plaque. First, we used CEA specimens and applied a highly specific LC-MSMS method to directly determine the chlorinating activity of hypochlorous acid that is specific for MPO.^25^ This method has been validated for the determination of MPO activity in atherosclerotic plaque.^23^ Using this method, we show that MPO activity is increased selectively in AHA type VI lesions. Second, MPO activity determined by *ex vivo* MPO-Gd retention and based on T1 mapping showed that R1 relaxation values were significantly higher in ruptured compared with stable plaques, the phenotype of which was confirmed by histology. Third, *ex vivo* MPO-Gd retention correlated with AHA plaque classification on *in vivo* carotid MRI, as well as MPO localization and vulnerable plaque features such as IPH determined by histology and IHC. As the paramagnetic effects of methemoglobin present in IPH increase R1 values, a pre-contrast baseline with the same T1 mapping sequence will be required for the detection of changes in R1 values driven by MPO-Gd accumulation in the presence of IPH. As T1 mapping protocols are already used routinely in clinical cardiovascular applications,^29^ an imaging strategy involving pre- and post-contrast T1 maps can be implemented readily in future *in vivo* translational studies.

Previous studies reported circulating concentrations of MPO protein to associate with atherosclerotic plaque instability (cardiovascular event risk),^30, 31^ and plaque erosion.^32, 33^ However, other studies showed no link between circulating MPO protein and cardiovascular risk.^34, 35^ Most studies demonstrating a correlation between circulating MPO protein and future events used enriched populations who had already experienced plaque destabilization. Like these studies, we observed plasma concentration of MPO protein to be elevated in patients with ruptured compared with stable plaques (**Figure S8B**). Interestingly however, circulating MPO protein concentrations were comparable in rabbits irrespective of future plaque destabilization (**Figure S8A**), despite elevated MPO activity being seen in animals that proceeded to atherothrombosis. This implies that MPO activity detected by molecular MRI may be a more sensitive and specific biomarker than plasma MPO concentrations. Irrespective of our findings, measuring circulating MPO protein has several limitations. Firstly, the absence of an approved universal protocol for quantifying plasma MPO protein limits its clinical use and the ability to compare absolute concentrations between studies. Secondly, determining plasma MPO protein does not provide insight into the chlorinating activity of the enzyme. Thirdly, it remains to be established whether MPO protein provides an incremental value to established plasma biomarkers for cardiovascular risk prediction such as CRP. Fourthly, plasma MPO protein is not causally associated with cardiovascular mortality.^36^ Lastly, plasma MPO concentrations provide no spatial information of plaques that may be susceptible to destabilization to guide intervention. These limitations can be overcome by measuring plaque MPO activity using molecular MRI with the apparent advantage of being able to predict plaque disruption prior to an event.

There has been continued effort to establish non-invasive imaging strategies that reliably detect vulnerable plaque based on multi-sequence MRI characterization of features of high-risk plaque^37, 38^ and disease activity using molecular imaging.^39, 40^ Such an approach can guide optimal therapy through prevention and intervention strategies for acute events precipitated by atherosclerotic plaque disruption and may improve clinical outcomes.^40, 41^ Plaque enhancement with non-selective contrast agents and high-resolution MRI associates with acute cardiovascular events,^42^ and the dynamics of the contrast agent uptake correlates with plaque inflammation.^43^ Nevertheless, enhancement reveals no further information about plaque activity due to the lack of specificity of contrast agents, which can be influenced by other factors that drive contrast uptake, such plaque vascularity. As a result, there has been ongoing interest in detecting specific inflammatory activity to denote plaque vulnerability as this could improve risk stratification and outcomes.^44, 45^

Current research strategies using positron emission tomography (PET) for imaging atherosclerotic plaque inflammation have focused on probes including ^18^F-fluorodeoxyglucose (FDG) and ^68^Ga-DOTATATE. The widely studied FDG-PET targets enhanced glucose uptake, an indirect proxy of general inflammation-related plaque metabolism. Though FDG-PET can detect plaque inflammation driven by the accumulation of macrophages, it is non-specific,^46^ susceptible to changes in blood glucose concentrations,^47^ generates signals with high heterogeneity,^48^ and cannot differentiate M1/M2 subsets of polarized macrophages which play juxtaposing roles in the inflammatory cascade. ^68^Ga-DOTATATE targets more specifically the somatostatin receptor subtype-2 but is limited by its inability to distinguish beneficial from harmful inflammation.^49, 50^ Alternatively, ultra-small superparamagnetic iron oxide (USPIO), a clinically approved MRI contrast agent, can track and characterize macrophage-infiltration.^51, 52^ However, unlike Gd-based MRI agents that can be imaged within 1 h of administration, the typically 12-24 h time gap between pre- and post-contrast injection MRI acquisitions required for USPIO is impractical clinically and may preclude the possibility of short-term follow-up with other diagnostic MRI sequences.^51^ USPIO may also preferentially label the anti-inflammatory M2-polarized macrophages,^53^ making it less suitable of identifying harmful inflammation.

Given the causal and specific role of extracellular MPO in plaque destabilization and vascular inflammation,^23, 24, 54^ MPO is a molecular target of interest in this emerging field offering a specificity to the disease process beyond current probes that target inflammation more broadly and/or indirectly.^17^ Enhancement with the MPO-activatable MRI agent has been correlated with active cardiovascular inflammatory lesions such as ischemic injury of the myocardium,^21^ and atherosclerotic plaques in different animal models.^19, 23^ MPO-Gd enhanced MRI provides selective and direct information on detrimental plaque inflammation induced by extracellular MPO. It detects pathologic plaque activity with high sensitivity and resolution and can be coupled with established multi-contrast MRI sequences which characterize high-risk plaque components,^37, 38^ providing an enhanced understanding of plaque vulnerability and more rapid and accurate risk stratification. The present study demonstrates this synergy and further specifies the capacity of imaging MPO activity to identify thrombosis-prone plaques and culprit lesions. Imaging plaque MPO activity using MPO-Gd MRI also has the advantage that the imaging can directly guide delivery and monitor the effect of existing investigational MPO inhibitors.^55^ Therefore, MPO activity may provide a coupled diagnostic and therapeutic strategy to deliver personalised treatment to maximize patient care.

MPO-Gd appears to be a realistic candidate for translational use. It has good biocompatibility^22^ as its main chemical components naturally occur in humans (5-hydroxytryptamine) or are already used in clinical imaging (DTPA-Gd). The imaging (30-90 min post-injection) and injected dose (0.1 mmol/kg, comparable to that used in patients) makes this approach suitable and practical for clinical application. Moreover, *in vivo* MRI in the present study was conducted on a clinical scanner using vessel wall imaging protocols and quantitative T1 mapping sequences commonly used in patients. These parameters render MPO-Gd the potential to aid the clinical detection of vulnerable plaques for treatment escalation, intervention, and surveillance. Further supporting this, a newer generation highly efficient MPO-activatable MRI probe, in which the linear DTPA chelator is replaced with a macrocyclic DOTA, has being developed.^56^ This may prove a superior candidate for human use given the improved gadolinium stability and faster blood clearance, eliminating potential issues of Gd toxicity, with background signal and shortening of the time between pre- and post-contrast imaging. Importantly, recent technical advancements in MRI employing accelerated image acquisition protocols, magnetic resonance fingerprinting, motion correction and deep-learning based reconstruction may further help to reduce MRI scan time and overcome current limitations of MRI to image atherosclerosis in patients.^8, 57, 58^ Moreover, advancements in software and hardware design have led to an improvement in signal-to-noise ratio for low-field MRI scanners (e.g., 0.55T). As Gd-based agents have higher R1 relaxivity at lower field strengths, it generates greater signal enhancement than at higher field strengths. This may facilitate future translation of molecular MRI strategies by decreasing the injected dose of Gd-based imaging probes and related costs. Considering the regulatory challenges for translating MRI-Gd probes, an alternative route to the clinical translation of imaging MPO activity may be the use of analogue MPO PET tracers,^59^ which a hybrid PET/MRI approach can take advantage of measuring the molecular imaging signal by PET and multi-sequence plaque characterization by MRI.

There are several limitations to this study. Firstly, in the rabbit model used, plaque disruption does not happen spontaneously but required exogenous triggers, although spontaneous atherothrombosis is very scarce in commonly used animal models of atherosclerosis.^60^ Secondly, the rabbit model lacks developing plaque calcification common in human plaques, and IPH which associates with lesion instability. Although both features have been previously reported in plaques of rabbits that were fed with a cholesterol-fortified diet for prolonged periods, animal mortality was increased. Other specific interventions to induce atherothrombosis in animal models that develop more advanced plaques are hampered by low incidence of atherothrombosis.^60^ Thirdly, due to the nature of human tissue collection, it was not possible to assess whether MPO activity predicts plaque rupture in humans, though the rabbit model data were specifically utilized to support this hypothesis. Finally, it was not possible to assess MPO activity in human tissue *in vivo*. As a result, the assessment of the probe in this study was limited to *ex vivo* soaking, probe activation and washing of CEA specimens. Notwithstanding these limitations, the preclinical and clinical parts of the present study are complementary and strengthen the case for future prospective clinical trials.

In conclusion, *in vivo* imaging of MPO activity using an MPO-Gd probe predicts future atherothrombosis in a preclinical model, and MPO activity is increased in ruptured human atherosclerotic plaque. These results highlight the unique and specific role of MPO activity and its potential application as a molecular target in the detection of vulnerable atherosclerotic plaque.

## Acknowledgments

We acknowledge the support of the Heart Research Institute mass spectrometry facility, and the University of New South Wales Biological Resource Imaging Laboratory.

## Non-standard Abbreviations and Acronyms

2-Cl-E^+^: 2-chloroethidium
AHA: American Heart Association
AWA: aortic wall area
CEA: carotid endarterectomy
DOTA: gadolinium-gadoteric acid
DTPA: diethylenetriaminepentaacetic acid
FDG: ^18^F-fluorodeoxyglucose
IPH: intraplaque hemorrhage
LC-MSMS: liquid chromatography tandem mass spectrometry
LGE: late MPO-Gd-enhanced
MPO: myeloperoxidase
MPO-Gd: gadolinium-*bis*-5-hydroxytryptamide diethylenetriaminepentaacetic acid
MRI: magnetic resonance imaging
PET: positron emission tomography
R1: relaxation rate
ROC: receiver operating characteristic
T1BB: 2D T1-weighted black blood
T1w-IR: 3D inversion recovery T1w
TMCR: tissue-to-muscle contrast ratio
USPIO: ultra-small superparamagnetic iron oxide

